# Myrmecophytism as a driver of macroevolutionary patterns: perspectives from the Southeast Asian *Macaranga* ant-plant symbiosis

**DOI:** 10.1101/2023.12.12.571248

**Authors:** Nadi M Dixit, Daniela Guicking

## Abstract

Myrmecophytes utilise defensive services offered by obligate ant partners in a novel means of survival in tropical habitats. Although much is known about the ecology of myrmecophytism, there aren’t enough empirical examples to demonstrate whether it substantially influences evolutionary patterns in host plant lineages. In this study, we make use of the species-rich *Macaranga* (Euphorbiaceae) ant-plant symbiosis distributed in Sundaland to delve into the evolutionary dynamics of myrmecophytism in host plants. We generated the most comprehensive dated phylogeny of myrmecophytic *Macaranga* till date using sequences derived from genotyping-by-sequencing (GBS), a next-generation sequencing (NGS) technique. With this in hand, we traced the evolutionary history of myrmecophytism in *Macaranga* using parametric biogeography and ancestral state reconstruction. Diversification rate analysis methods were employed to determine if myrmecophytism enhanced diversification rates in the genus. Our results demonstrate that myrmecophytism is plastic and easily lost unless it is overly specialised. Ancestral state reconstruction supported a single origin of myrmecophytism in *Macaranga* ∼18 mya on Borneo followed by multiple losses. Diversification rate analysis methods did not yield sufficient evidence to support the hypothesis that myrmecophytism enhanced diversification rates in *Macaranga*; we found that topographical features on Borneo may have played a more direct role in the divergence of clades instead. Through this comprehensive investigation of the evolution of myrmecophytism in *Macaranga*, our study also provides evidence that a key innovation may not necessarily enhance diversification rates. In fact, we hypothesise that overly specialised cases of myrmecophytism may even be an evolutionary dead end.

## 1. Introduction

Key innovations are defined as traits that allow organisms to interact with their environment in novel ways (Miller, 1949; Liem, 1973; Losos & Mahler, 2010) and thereby enable them to enter into a previously inaccessible ecological state (Miller *et al*., 2022). The acquisition of such novel traits can promote differential evolutionary success among lineages (Larouche *et al*., 2020) through the creation of new adaptive zones, access to previously unattainable resources, and increased fitness (Losos & Mahler, 2010; Fernández-Mazuecos *et al*., 2019). The concept of a key innovation has been rather controversial in recent years with multiple studies incorporating increased lineage diversification as a core aspect of its definition. Others argue that speciation could be hindered by factors such as high levels of gene flow or even a lack of evolvability in spite of a significant ecological shift (Vrba, 1987; Schluter, 2000; Claramunt *et al*., 2012; Payne & Wagner, 2019; reviewed in Miller *et al*., 2022), implying enhanced diversification isn’t necessarily a consequence of a key innovation. Hence, in an attempt to recognise the two as distinct phenomena, Miller *et al*. (2022) suggest an additional term, “diversifying trait”, to specifically refer to those traits whose evolution leads to increased diversification rates in a clade. There are, however, traits known to fall under both categories (Miller *et al*., 2022).

Nectar spurs, tubular outgrowths of a floral organ that usually contain nectar, are considered a classic example of a plant key innovation that promoted diversification in angiosperms by means of pollinator shifts (Hodges & Arnold, 1995; Whittall & Hodges, 2007; Puzey *et al*., 2012; Fernández-Mazuecos *et al*., 2019). Similarly, unique plant traits whose evolution has triggered generalist mutualistic interactions with ants qualify as both key innovations and diversifying traits. These include extrafloral nectaries (EFNs), nectar secreting glands found on non-floral plant tissues that provide trophic incentives to ants in return for defensive services (Bentley, 1977), and elaiosomes, nutritive fatty appendages on seeds meant for ant dispersers (Berg, 1975). Both traits have allowed plants to make an ecological shift in their means of survival and reproduction (Heil & McKey, 2003; Bronstein *et al*., 2006; Lengyel *et al*., 2010), while also influencing macroevolutionary patterns in multiple plant lineages by enhancing diversification rates (Lengyel *et al*., 2009; Weber & Agrawal, 2014). However, it is still not clear whether the more specialised and obligate variant of ant-plant mutualism, myrmecophytism, similarly qualifies as both. In this almost exclusively tropical interaction (Chomicki & Renner, 2015), host plants, so-called myrmecophytes, provide substantially more valuable rewards to their ant partners than their generalist counterparts: specialised nesting spaces, called domatia, in hollow stem internodes, thorns, petioles or leaf pouches, in addition to trophic resources, usually in the form of nutrient-rich food bodies. Together these rewards ensure a very close association between plants and ants (Heil *et al.,* 2001; Heil & McKey, 2003; Bronstein *et al*., 2006). The nesting ants show high fidelity to their host plants and protect them with great efficiency against herbivory, competition from encroaching vines, and even from pathogens (Davidson & McKey, 1993; Heil & McKey, 2003; Bronstein *et al*., 2006; Rico-Gray & Oliveira, 2007). Field studies have demonstrated that myrmecophytic host plants without their ant partners show significant leaf damage and are usually overgrown by competitive vines (Janzen, 1966; Heil & McKey, 2003), indicating their lack of survivability without their ant partners. In some epiphytic myrmecophytes, host plants also make use of ant debris such as excrement, dead larvae and workers deposited in their domatia as a source of nutrition (Beattie, 1989; Chomicki & Renner, 2017). Clearly, this novel defensive and, in some cases, nutritional regime has enabled host plants access to a significant and effective means of survival in habitats where combating herbivory, competition or unreliable nutrient supply becomes integral to survival. Given these observations, myrmecophytic associations can be considered a key innovation that represents a unique adaptation in tropical plants to counter elements unfavourable to viability.

Unlike the more generalised interactions, myrmecophytism is a relatively rare phenomenon and is known to occur in only around 700 species from across 50 families of vascular plants (Chomicki & Renner, 2015; Nelsen *et al*., 2018). In contrast, EFNs are found in 4,000 species from 100 angiosperm and fern families, and elaiosomes are known from around 11,000 species across 77 angiosperm families (Nelsen *et al*., 2018). This raises the question as to whether or not myrmecophytism is also a diversifying feature. In their study on the global scatter of domatium evolution in vascular plants, Chomicki & Renner (2015) suggested that the formation of domatia, in general, did not have a significant impact on diversification rates in host plant lineages. Their relatively rare prevalence, frequent association with species-poor clades, and scattered phylogenetic pattern of occurrence in fact suggest that domatia could be easily lost or that myrmecophytism might even represent an evolutionary dead-end (Peccoud *et al*., 2013; Chomicki & Renner, 2015). Nonetheless, a few species-rich myrmecophytic plant genera such as *Neonauclea*, *Cecropia,* and *Macaranga* exist and it could be possible to hypothesise myrmecophytism-mediated increased diversification rates at least in these cases (Blattner *et al*., 2001; Weising *et al*., 2010; Chomicki & Renner, 2015).

The *Macaranga*-*Crematogaster* symbiosis is among the most species-rich and best-studied ant-plant systems worldwide, making it an ideal system for studying the evolution of myrmecophytism. *Macaranga* belongs to the Euphorbiaceae family and comprises ∼300 paleotropically distributed tree and shrub species, many of which inhabit early successional habitats (Davies, 2001; Whitmore *et al*., 2008). While many *Macaranga* species facultatively attract ants by offering EFNs and/or food bodies, ∼30 species distributed in the Southeast Asian tropics, specifically Sundaland, participate in myrmecophytic associations with primarily nine species of *Crematogaster* ants which nest in the hollow internodes of their host plants (**Fig**. **1**, Supplementary material **Table S3**; Fiala *et al*., 1989, 1999; Heil *et al*., 2001; Quek *et al*., 2007; Feldhaar *et al*., 2010, 2016). Besides nesting spaces, the host plants offer nutrition in the form of lipid-rich food bodies to their ant partners (Fiala & Maschwitz, 1992b; Fiala *et al*., 1999). In most *Macaranga*-associated partnerships, the ants additionally tap nutrients from honeydew derived from scale insects (family Coccidae) that they rear inside the domatia of their host plants (Heckroth *et al*., 1998; Gullan *et al*., 2018). EFNs are usually also present in *Macaranga* myrmecophytes but are highly reduced and apparently don’t play a major role in feeding the ant partners (Fiala & Maschwitz, 1991; Fiala *et al*., 1999). Under natural conditions, the myrmecophytes cannot survive to reproductive age without their ant partners and the *Crematogaster* ant partners have so far only been found on their *Macaranga* host plants, signifying the obligate symbiotic nature of this association (Fiala *et al*., 1989; Heil *et al*., 2001).

**Fig. 1.**
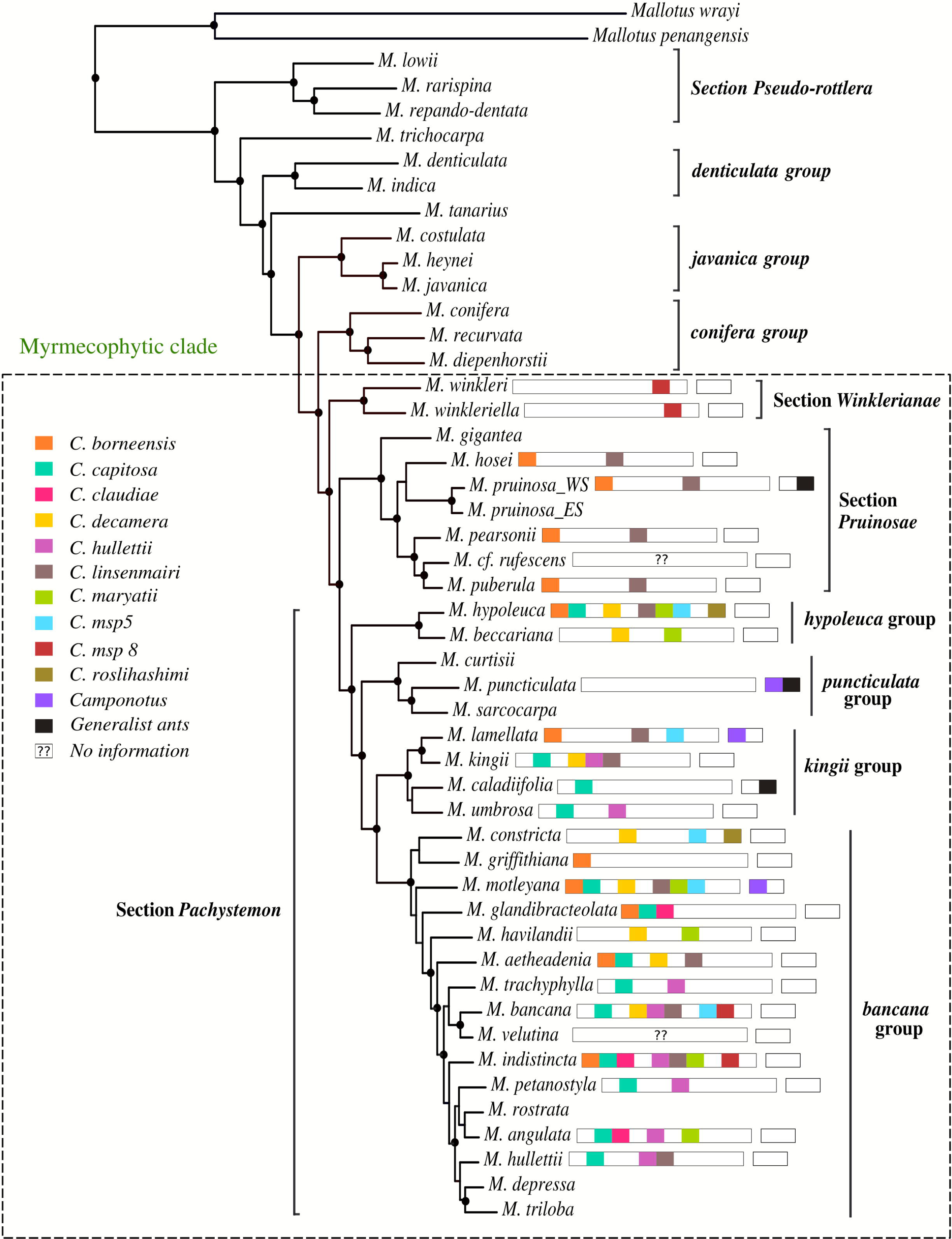
Phylogenetic tree and specificity of the ant-plant interaction. RAxML maximum likelihood tree estimation of 47 *Macaranga* taxa along with two *Mallotus* species functioning as outgroups. Nodes that received bootstrap support > 95% are marked with black solid circles. The clade comprising all three myrmecophytic taxonomic sections: *Pachystemon*, *Pruinosae*, and *Winklerianae* has been marked with a dashed square and indicated as the myrmecophytic clade. Two individuals representing the myrmecophytic and non-myrmecophytic forms of *M. pruinosae* are denoted by their distinct geographic distributions: WS (West Sundaland, comprising Sumatra and Malay Peninsula) and ES (East Sundaland, comprising Borneo) respectively. Within the myrmecophytic clade, partner ants of each myrmecophytic species are indicated in bars as coloured blocks next to the corresponding host plant species. Primary ant partners of the *Macaranga* myrmecophytes that belong to the *Crematogaster* genus are indicated in the longer block on the left while less frequent partnerships with *Camponotus* species and/or generalist ants (**Table S3**) are indicated in the smaller block on the right. Ant species are colour coded as illustrated in the legend. No information on ant partner species is yet available for *M. cf. rufescens* and *M. velutina*. *Crematogaster msp5* and *C. msp8* are morphospecies which have not been formally described yet. Information on ant partners of the myrmecophytes was gathered from Fiala (1999) and Feldhaar *et al*. (2016).

Previous attempts to investigate the evolution of myrmecophytism in *Macaranga* suffered from shortcomings such as poor resolution of interspecific relationships, uncertain species identities and designations, and insufficient sampling (Blattner *et al*., 2001; Davies *et al*., 2001; Bänfer *et al*., 2004; Kulju *et al*., 2007; Chomicki & Renner, 2015; Fiala *et al*., 2016). Nevertheless, it has been hypothesised that the acquisition of obligate ant partners in *Macaranga* may have accelerated diversification by enhancing competitiveness of host plant lineages in pioneer habitats (Blattner *et al*., 2001; Weising *et al*., 2010). To investigate the impact of an evolutionary innovation on diversification, it is not possible to single it out as a stand-alone causal factor, as ecological context is an important determinant in evolution (Hunter, 1998). In this regard, the effect of abiotic and ecological factors on the diversity of the taxa, in addition to stochasticity, cannot be excluded (Cracraft, 1990). Numerous studies have shown the influence of multiple, sometimes interdependent, ecological and abiotic factors on mutualistic interactions, e.g. habitat type, altitude, and geological processes such as mountain building (Lagomarsino *et al*., 2016; Chomicki & Renner, 2017; Jimenez *et al*., 2021). In symbiotic associations, the abundance of mutualistic partners can vary across habitats and factors such as resource availability and interspecific competition can shift cost-benefit ratios to either promote mutualistic associations or cause them to break down.

In *Macaranga*, the relatively large number of myrmecophytic species with diverse ecologies, the prevalence of varied degrees of specialisation with respect to ant-partners, the distribution of host plants in a number of different habitat types, and the geological complexity of Sundaland make it an ideal system to gain deeper insights into the evolution of myrmecophytism and its impact on the evolutionary dynamics of the host-plant lineages. To accomplish this, we employ genotyping-by-sequencing (GBS; Elshire *et al*., 2011), a next-generation DNA sequencing (NGS) technique, to generate the most comprehensive and well-sampled dated phylogeny of myrmecophytic *Macaranga* and its closest non-myrmecophytic relatives to date. With this framework, we specifically address the following questions (1) When and where did myrmecophytism evolve in *Macaranga* and what were the conditions that promoted its origin? Have there been multiple instances of origins and losses of myrmecophytism in *Macaranga* and in what ecological contexts did they occur? (2) Did myrmecophytism enhance diversification rates in *Macaranga*? (3) Are there any characteristic evolutionary patterns in *Macaranga* host plant lineages that could be associated with the acquisition of obligate ant partners?

## 2. Materials and Methods

### 2.1. Taxon sampling and DNA sequence dataset

Our sampling includes representatives from all taxonomic subdivisions of myrmecophytic *Macaranga* and their closest non-myrmecophytic *Macaranga* lineages distributed in Sundaland. Myrmecophytic *Macaranga* species occur in three taxonomic sections: *Pachystemon*, *Pruinosae*, and *Winklerianae* (Davies, 2001; Whitmore *et al*., 2008). Section *Pachystemon* is further divided into four subgroups: *bancana*, *hypoleuca*, *kingii*, and *puncticulata* based on phylogenetic analyses (Bänfer *et al*., 2004). We included a total of 136 individuals representing 33 species from the three myrmecophytic sections and an additional 13 species from the non-myrmecophytic *Macaranga* lineages (*conifera* group, *denticulata* group, *javanica* group, section *Pseudo-rottlera*, *tanarius* group, and *trichocarpa* group) to generate a comprehensive phylogeny (Supplementary material, **Fig. S1**). From this larger sampling, a subset comprising a single representative of each of the 46 species included here (with the exception of two individuals from *Macaranga pruinosa*, representing both its myrmecophytic and non-myrmecophytic forms) was used to trace evolutionary patterns. Two individuals from *Mallotus*, sister genus to *Macaranga* (Kulju *et al*., 2007; Sierra *et al*., 2010; van Welzen *et al*., 2014), were included as outgroups. All information on methodology and results pertaining to the larger sample set is detailed in Supplementary material (**Fig. S1**)

Sourcing of plant material, DNA extraction, GBS library preparation, sequence alignment and SNP mining protocols followed Dixit *et al*. (2023), except for the sequence clustering threshold parameter in the ipyrad assembly pipeline which was set at 90%. The resulting sequence assembly had 2,097,049 sites for 49 individuals with 50.76% missing sites. The SNP matrix had 212,516 sites and the unlinked SNP matrix had 14,710 sites in total.

### 2.2. Maximum likelihood phylogenetic analysis

Maximum likelihood (ML) phylogenetic tree calculation was performed using the ipa.raxml() tool available in the ipyrad analysis toolkit (Eaton & Overcast, 2020). The tool automates the process of generating RAxML (Stamatakis, 2014) command line strings and running them through Python code. For our data, the generalized time reversible (GTR) model was chosen as the substitution model along with the GAMMA model for rate heterogeneity (GTRGAMMA). A rapid bootstrap analysis with 100 replicates and the search for the best-scoring ML tree were simultaneously conducted in one single program run. The tree was rooted with two *Mallotus* individuals: *Mallotus penangensis* and *M. wrayi*. Toytree (Eaton, 2020) was used to plot and visualise the best-scoring tree.

### 2.3. Divergence time estimation and historical biogeographic analysis

Bayesian phylogenetic inference along with divergence time estimation was performed on BEAST2 (version 2.6.3; Bouckaert *et al*., 2019) using the unlinked SNP matrix obtained from the ipyrad assembly pipeline. The bModelTest package (Bouckaert & Drummond, 2017) was implemented through model averaging to determine the site model. A relaxed uncorrelated lognormal clock was used and speciation was set to occur according to a Yule process (Drummond *et al*., 2006). Implementing the Birth-Death model showed negligible differences in tree topology and dates in comparison to the Yule model runs. Four dated nodes derived from van Welzen *et al*. (2014) served as secondary calibration points and were coded as MRCA uniform distribution priors with limits derived from the 95% HPD intervals of the corresponding secondary calibration estimates (Ho, 2007; Kuriyama *et al*., 2011; Ryberg & Matheny, 2011; Siler *et al*., 2012; Kornilios *et al*., 2013; Ruiz-Sanchez & Specht, 2013; Schenk, 2013; Zhao *et al*., 2013; Schenk, 2016): crown node of the *Macaranga-Mallotus* clades (95% HPD: 63.33-79.13 mya), stem node of section *Pseudo-rottlera* (95% HPD: 28.01-41.46 mya) *M. gigantea-M. pearsonii* split (95% HPD: 2.17-9.46 mya), *M. umbrosa-M. lamellata* split (95% HPD: 0.64-5.78 mya). The Markov Chain Monte Carlo (MCMC) chain length was set to 50,000,000 generations from which every 2000th generation was sampled. Convergence of the chain was monitored on Tracer (version 1.7.1; Rambaut *et al*., 2018) and ESS>200 was used as a criterion to check for the quality of the resulting sample sequence. The maximum clade credibility tree from among the sampled trees was annotated using TreeAnnotator (version 2.6.3 BEAST2 package) and visualised on FigTree (version 1.4.4; Rambaut, 2019).

To infer biogeographic origins of myrmecophytism in *Macaranga*, ancestral area reconstruction was performed on RASP (version 4.0.0, Yu *et al*., 2020). The geographic ranges of the extant taxa were divided and coded as follows: A = Indian subcontinent + Indochina + South China + Japan, B = Malay + Thai Peninsulas, C = Sumatra, D = Java, E = Borneo, F = Philippines, G = Sulawesi + Maluku islands + New Guinea + Australia and all the minor islands in east Pacific. Since our study focusses on the myrmecophytic species that are all restricted to Sundaland, the very broad distribution ranges of non-myrmecophytes that fall outside of Sundaland and thus not relevant to our study questions were compressed into the single areas A, F, and G. In addition, a higher number of areas resulted in a very large number of ancestral area combinations that significantly increased computation time. Hence, bringing down the total number of areas to seven by combining geographic ranges was deemed optimal. The reconstruction of ancestral ranges was performed by implementing the six BioGeoBEARS models (DEC, DIVALIKE, BAYAREALIKE and their +j counterparts; Matzke, 2014) on RASP. The best estimation was chosen by comparing the likelihood and AIC values of the six models.

### 2.4. Ancestral state reconstruction

The evolutionary history of myrmecophytism in *Macaranga* was inferred through ancestral state reconstruction of this trait (treated as a single character) on Mesquite (version 3.70 Maddison & Maddison, 2021). The reconstruction was carried out by using likelihood (Mk1 equal transition rates and AsymmMk unequal transition rates models) and parsimony methods. These were implemented on the maximum clade credibility (MCC) tree derived from BEAST2 and on the best-scoring ML tree calculated with the RAxML tool to account for phylogenetic discrepancies between the two estimates. Likelihood ratio test was used to compare the two likelihood models.

### 2.5. Estimation of diversification rates

In order to test whether the acquisition of myrmecophytism noticeably spurred diversification rates in *Macaranga*, both state-dependent and state-independent methods were employed. The hypothesis that myrmecophytism enhanced diversification rates in *Macaranga* was specially raised on the background of high species diversity in the *bancana* group of section *Pachystemon* (Blattner *et al*., 2001; Weising *et al*., 2010). Although this group has been considered “hyperdiverse” based on species descriptions in literature (Blattner *et al*., 2001; Davies, 2001; Weising *et al*., 2010), our preliminary analysis of GBS data suggests that the species number in this group may have been overestimated in earlier work (Supplementary material **Fig. S3**). Since the number of species included in a clade is known to affect the outcome of diversification rate analyses (Faurby *et al*., 2016), we decided to account for this taxonomic uncertainty in our analyses. Hence, following the approach of Fernández-Mazuecos *et al*. (2019), we specify two alternative taxonomic treatments of the *bancana* group: “splitter” and “lumper“. In the “splitter” treatment all 16 individuals that are currently recognized as distinct species in the *bancana* group in the latest taxonomic revision of section *Pachystemon* (Davies, 2001) were included. In the “lumper” treatment, the total number of species was reduced from 16 to 8 following the results of our preliminary STRUCTURE analysis (Supplementary material, **Fig. S3**) of the *bancana* group, including only one individual from each “distinct” cluster (Supplementary material **Fig. S3**). The diversification rate estimation methods were applied to both of these treatments.

#### 2.5.1. Trait-dependent methods

Various models under two trait-dependent methods were applied to identify diversification rate shifts in *Macaranga*: BiSSE (binary state speciation extinction; FitzJohn *et al*., 2009) and HiSSE (hidden state speciation extinction; Beaulieu & O’Meara, 2016).

BiSSE estimates diversification rates associated with a binary trait in a model-based approach. BiSSE was implemented in the R package diversitree (FitzJohn, 2012) on the MCC tree obtained from BEAST2 for both the splitter and lumper taxonomic treatments. Incomplete sampling was accounted for by including proportions of non-myrmecophytes and myrmecophytes in *Macaranga* that were not included in this study. Eight models, including the full BiSSE model, were specified by constraining various combinations of speciation rate (A), extinction rate (µ) and transition rate (q) parameters to be equal (**Table 1a**); parameters were estimated under each model in the ML approach and the best-fit model was chosen based on AIC values. To obtain a probability distribution for the parameter estimates, a Bayesian BiSSE analysis was conducted using exponential priors on ML parameter estimates of the full BiSSE model. The MCMC chain was set to run for 10,000 generations. The posterior probability distributions of the rates and their 95% credible intervals were then visualised as plots.

**Table 1.**
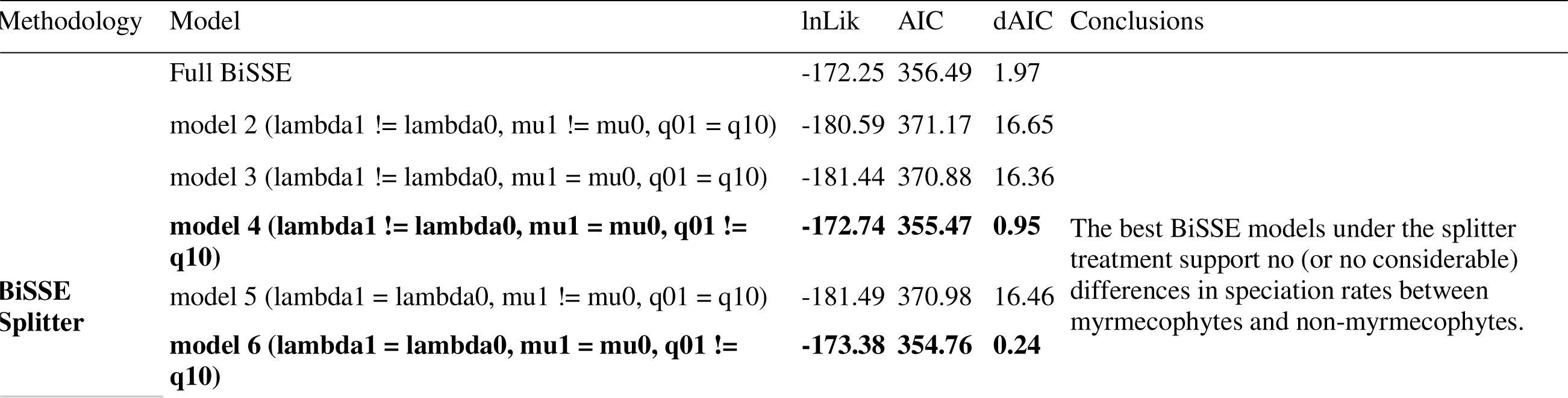
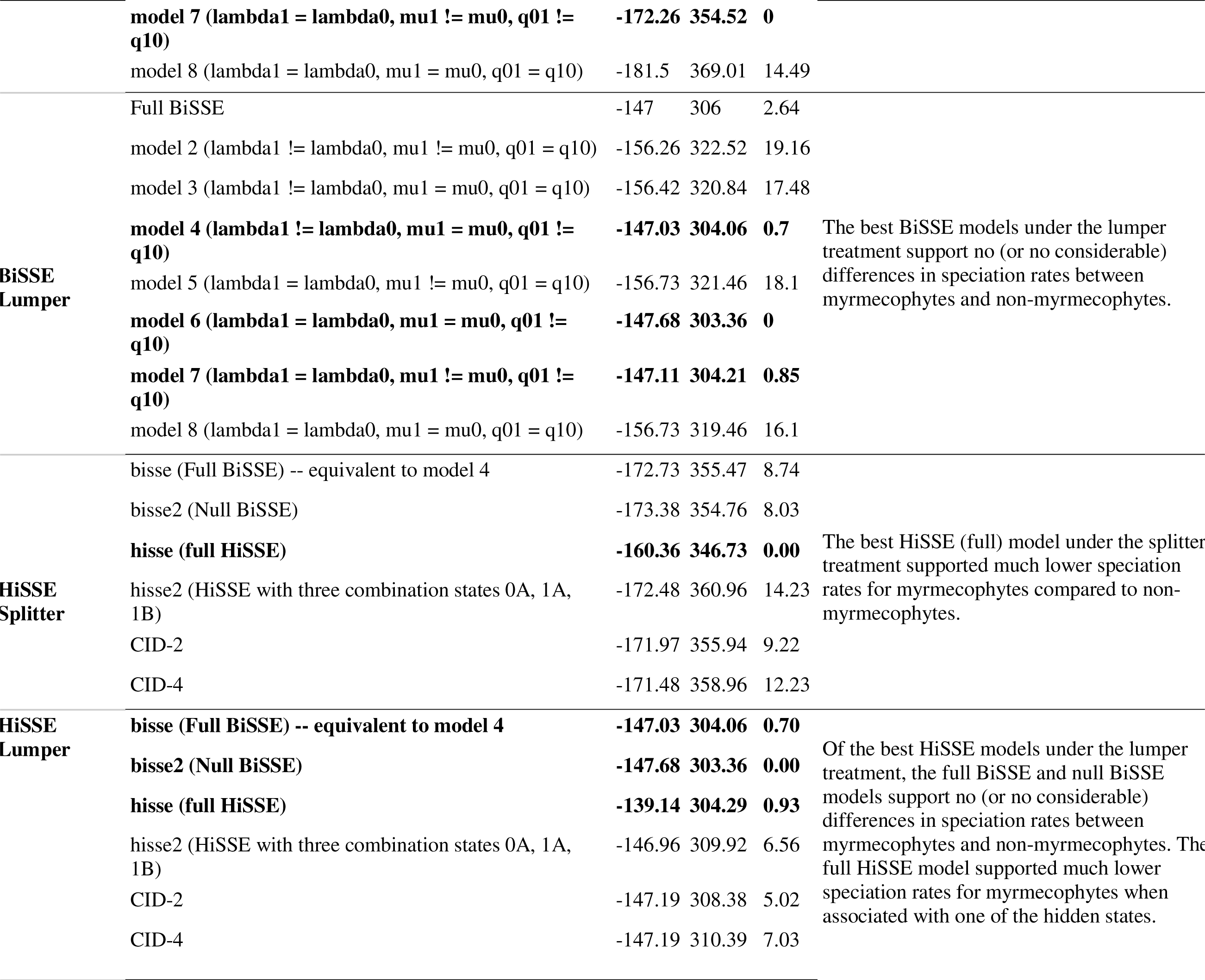
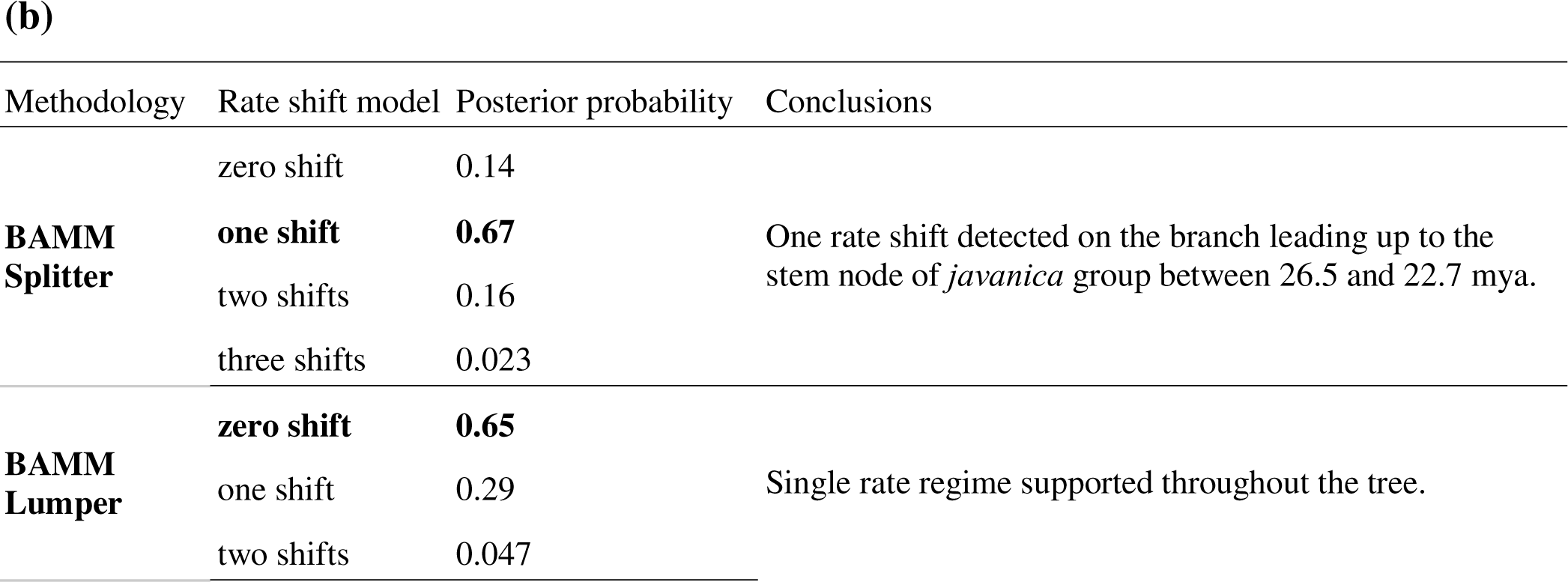
(a) Log-Likelihood and AIC values for various diversification rate analysis BiSSE (diversitree) and HiSSE (hisse) models estimated for both the splitter and lumper treatments. (b) Posterior probabilities for various rate shift models under the trait-independent BAMM analysis, implemented for both splitter and lumper treatments.

| Methodology | Model | lnLik | AIC | dAIC | Conclusions |
| --- | --- | --- | --- | --- | --- |
| BiSSE Splitter | Full BiSSE | -172.25 | 356.49 | 1.97 | The best BiSSE models under the splitter treatment support no (or no considerable) differences in speciation rates between myrmecophytes and non-myrmecophytes. |
| | model 2 ($\lambda_1 \neq \lambda_0$, $\mu_1 \neq \mu_0$, $q_0 = q_1$) | -180.59 | 371.17 | 16.65 | |
| | model 3 ($\lambda_1 \neq \lambda_0$, $\mu_1 = \mu_0$, $q_0 = q_1$) | -181.44 | 370.88 | 16.36 | |
|  | **model 4 ($\lambda_1 \neq \lambda_0$, $\mu_1 = \mu_0$, $q_0 \neq q_1$)** | **-172.74** | **355.47** | **0.95** |  |
| | model 5 ($\lambda_1 = \lambda_0$, $\mu_1 \neq \mu_0$, $q_0 = q_1$) | -181.49 | 370.98 | 16.46 | |
|  | **model 6 ($\lambda_1 = \lambda_0$, $\mu_1 = \mu_0$, $q_0 \neq q_1$)** | **-173.38** | **354.76** | **0.24** |  |
|  | **model 7 ($\lambda_1 = \lambda_0$, $\mu_1 \neq \mu_0$, $q_1 \neq q_0$)** | **-172.26** | **354.52** | **0** |  |
| | model 8 ($\lambda_1 = \lambda_0$, $\mu_1 = \mu_0$, $q_1 = q_0$) | -181.5 | 369.01 | 14.49 | |
| **BiSSE**<br>**Lumper** | Full BiSSE | -147 | 306 | 2.64 |  |
| | model 2 ($\lambda_1 \neq \lambda_0$, $\mu_1 \neq \mu_0$, $q_1 = q_0$) | -156.26 | 322.52 | 19.16 | |
| | model 3 ($\lambda_1 \neq \lambda_0$, $\mu_1 = \mu_0$, $q_1 = q_0$) | -156.42 | 320.84 | 17.48 | |
|  | **model 4 ($\lambda_1 \neq \lambda_0$, $\mu_1 = \mu_0$, $q_1 \neq q_0$)** | **-147.03** | **304.06** | **0.7** | The best BiSSE models under the lumpertreatment support no (or no considerable) differences in speciation rates between myrmecophytes and non-myrmecophytes. |
| | model 5 ($\lambda_1 = \lambda_0$, $\mu_1 \neq \mu_0$, $q_1 = q_0$) | -156.73 | 321.46 | 18.1 | |
|  | **model 6 ($\lambda_1 = \lambda_0$, $\mu_1 = \mu_0$, $q_1 \neq q_0$)** | **-147.68** | **303.36** | **0** |  |
|  | **model 7 ($\lambda_1 = \lambda_0$, $\mu_1 \neq \mu_0$, $q_1 \neq q_0$)** | **-147.11** | **304.21** | **0.85** |  |
| | model 8 ($\lambda_1 = \lambda_0$, $\mu_1 = \mu_0$, $q_1 = q_0$) | -156.73 | 319.46 | 16.1 | |
| **HiSSE**<br>**Splitter** | bisse (Full BiSSE) -- equivalent to model 4 | -172.73 | 355.47 | 8.74 |  |
|  | bisse2 (Null BiSSE) | -173.38 | 354.76 | 8.03 |  |
|  | **hisse (full HiSSE)** | **-160.36** | **346.73** | **0.00** | The best HiSSE (full) model under the splitter treatment supported much lower speciation rates for myrmecophytes compared to non-myrmecophytes. |
|  | hisse2 (HiSSE with three combination states 0A, 1A, 1B) | -172.48 | 360.96 | 14.23 |  |
|  | CID-2 | -171.97 | 355.94 | 9.22 |  |
|  | CID-4 | -171.48 | 358.96 | 12.23 |  |
| **HiSSE**<br>**Lumper** | **bisse (Full BiSSE) -- equivalent to model 4** | **-147.03** | **304.06** | **0.70** | Of the best HiSSE models under the lumpertreatment, the full BiSSE and null BiSSE models support no (or no considerable) differences in speciation rates between myrmecophytes and non-myrmecophytes. The full HiSSE model supported much lower speciation rates for myrmecophytes when associated with one of the hidden states. |
|  | **bisse2 (Null BiSSE)** | **-147.68** | **303.36** | **0.00** |  |
|  | **hisse (full HiSSE)** | **-139.14** | **304.29** | **0.93** |  |
|  | hisse2 (HiSSE with three combination states 0A, 1A, 1B) | -146.96 | 309.92 | 6.56 |  |
|  | CID-2 | -147.19 | 308.38 | 5.02 |  |
|  | CID-4 | -147.19 | 310.39 | 7.03 |  |

(b)
| Methodology | Rate shift model | Posterior probability | Conclusions |
| --- | --- | --- | --- |
| BAMM Splitter | zero shift | 0.14 | One rate shift detected on the branch leading up to the stem node of *javanica* group between 26.5 and 22.7 mya. |
|  | one shift | 0.67 |  |
|  | two shifts | 0.16 |  |
|  | three shifts | 0.023 |  |
| BAMM Lumper | zero shift | 0.65 | Single rate regime supported throughout the tree. |
|  | one shift | 0.29 |  |
|  | two shifts | 0.047 |  |

HiSSE is an extension to BiSSE by accounting for “hidden” states that could be influencing the diversification rates along with the observed character of interest (here, myrmecophytism). HiSSE was implemented in the R package hisse (Beaulieu & O’Meara, 2016) on the MCC tree while considering incomplete sampling proportions like in the BiSSE approach. Overall six models were specified: a full BiSSE model, a null BiSSE model, a full HiSSE model, a HiSSE model (referred to as hisse2 here) which allows only one of the observed binary states to co-occur with both states (A and B) of a binary hidden trait (0A, 1A, and 1B), a character-independent diversification model with two hidden states (CID-2), and a character-independent diversification model with four hidden states (CID-4). Rate parameters under each of these models were estimated and the best model fit was chosen based on AIC values (**Table 1a**).

#### 2.5.2. Trait-independent method

A state-independent approach was employed using BAMM v.2.5.0 (Bayesian analysis of macroevolutionary mixtures; Rabosky, 2014) to test if there is a discernible shift in evolutionary rates associated with the divergence of the myrmecophytic lineages. BAMM detects and quantifies heterogeneity in evolutionary rates in a Bayesian framework. Prior values for speciation and extinction parameters were generated using the *setBAMMpriors* function in the R package BAMMtools (Rabosky *et al*., 2014). Four MCMC chains were run for 1×10^7^ generations with every 5000th generation sampled. Ten percent of the output was removed as burn-in and convergence was assessed with the R package coda (Plummer *et al*., 2006). Incomplete sampling proportions were accounted for. The best rate-shift model was detected based on posterior probabilities and the best rate-shift configuration was visualised as a phylorate plot. Evolutionary rate trends were visualised as clade-specific rate-through-time (RTT) plots.

## 3. Results

### 3.1. Maximum likelihood phylogenetic tree construction

The ML phylogenetic reconstruction yielded a highly resolved and strongly supported tree, making the intersectional and interspecific relationships in myrmecophytic *Macaranga* explicit for the very first time, also with respect to their closest non-myrmecophytic relatives (**Fig. 1**, see Supplementary material **Fig. S1** for details pertaining to the larger sample set). The three myrmecophytic taxonomic sections, *Pachystemon, Pruinosae*, and *Winklerianae* (Davies *et al*., 2001; Whitmore *et al*., 2008), together form a monophyletic clade (from here on referred to as the myrmecophytic clade) while appearing as monophyletic clades themselves. Within section *Pachystemon*, the four subgroups: *bancana, hypoleuca*, *kingii*, and *puncticulata* previously recognised by Davies *et al*. (2001) and Bänfer *et al*. (2004) are each clearly resolved as monophyletic clades with a 100% bootstrap support. While almost all interspecific relationships within *Pruinosae*, *Winklerianae*, and three out of four subgroups within *Pachystemon* receive strong support, species relationships in the *bancana* group are not as well-supported (65-90%) and remain somewhat ambiguous. This finding is concordant with results from previous investigations which especially struggled to infer interspecific relationships in this particular group (Blattner *et al*., 2001; Davies *et al*., 2001; Bänfer *et al*., 2004). The non-myrmecophytes included in the study are also resolved with strong support.

### 3.2. Divergence time estimation and historical biogeographic analysis

Bayesian phylogenetic inference agreed with the ML estimation with respect to the positioning and composition of all taxonomic sections and groups (**Fig. 2**). There were, however, a few species that showed a slightly different positioning, particularly in the *bancana* group, but these corresponded to nodes that received relatively low bootstrap support in the ML estimation.

**Fig. 2.**
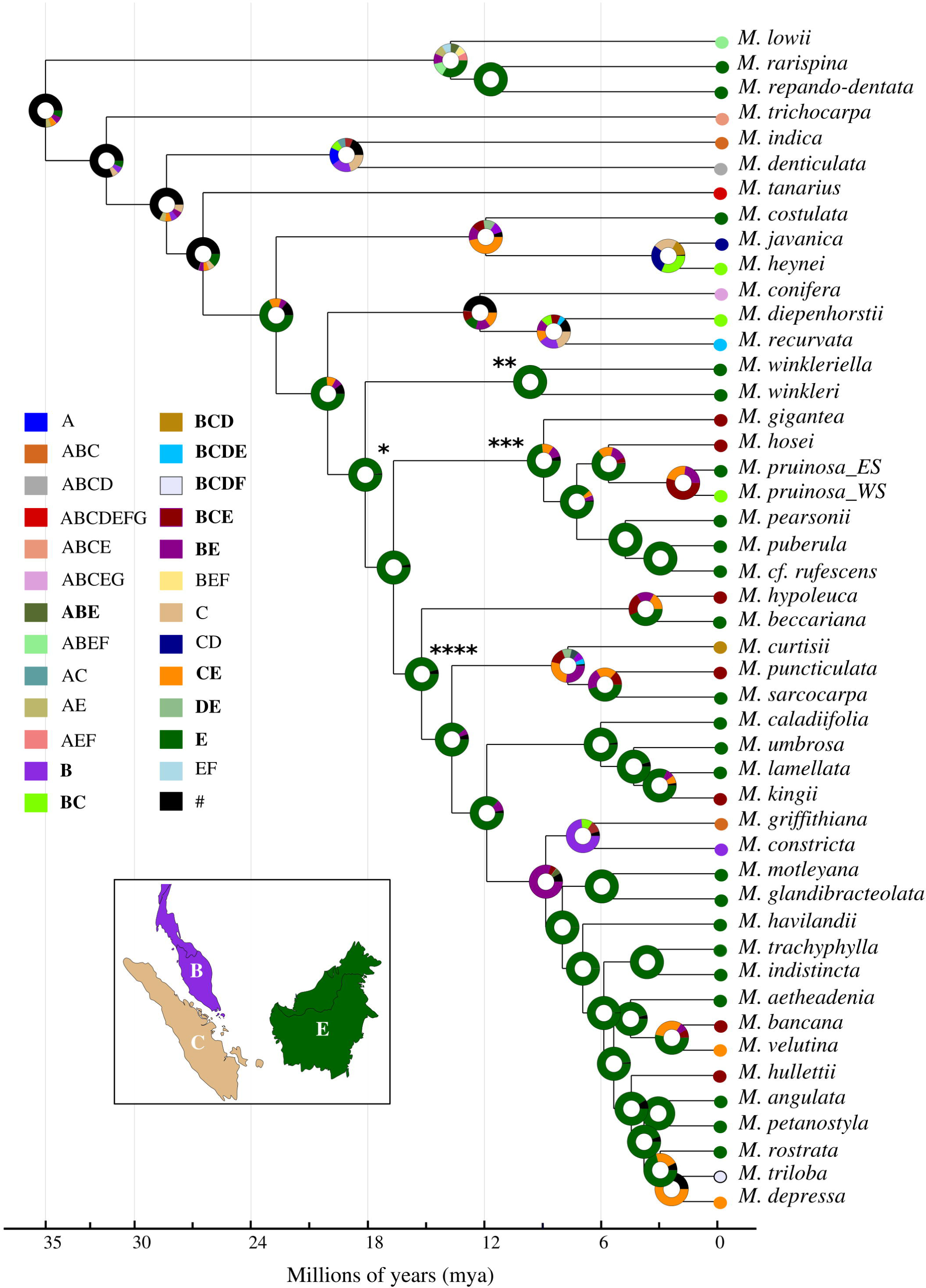
Divergence time estimation and historical biogeographic analysis. Ancestral area reconstruction of myrmecophytic *Macaranga* and their closest non-myrmecophytic relatives given by DEC model implemented on the time-calibrated maximum clade credibility tree estimated on BEAST2. Asterisks mark the crown nodes of the myrmecophytic clade (*), section *Winklerianae* (**), *Pruinosae* (***), and *Pachystemon* (****). The time scale at the base of the tree is calibrated in millions of years (mya) as indicated. Colour coded pie charts at each node represent distribution region or a combination of distribution regions occupied by ancestral taxa as estimated by the model (abbreviations: A = Indian subcontinent + Indochina + South China + Japan, B = Malay + Thai Peninsulas, C = Sumatra, D = Java, E = Borneo, F = Philippines, G = Sulawesi + Maluku islands + New Guinea + Australia and all the minor islands in east Pacific). The proportion of colours in the pie charts indicates the relative probability of a distribution region or combination of distribution regions to be the ancestral region. Black (#) is used to indicate the sum total of all regions or combination of regions that received probabilities < 5% in the estimation. Coloured circles at the tree tips show the present-day distributions of the respective taxa.

Specifically, the clade constituting *M. aetheadenia, M. angulata, M. bancana, M. depressa, M. hullettii, M. indistincta, M. petanostyla*, *M. rostrata, M. trachyphylla, M. triloba,* and *M. velutina* showed ambiguous interspecific relationships, which could point to incomplete lineage sorting.

The crown node of the myrmecophytic clade was dated at 18.13 mya (95% HPD: 16.38-23.6 mya) (**Fig. 2**, marked *), which falls in early-mid Miocene. The three myrmecophytic sections, *Winklerianae, Pruinosae* and *Pachystemon*, had their crown nodes dated at 9.64 mya (95% HPD: 4.83-14.43 mya) (**Fig. 2**, marked **), 8.95 mya (95% HPD: 8.03-9.49 mya) (**Fig. 2**, marked ***), and 15.22 mya (95% HPD: 12.35-18.06 mya) (**Fig. 2**, marked ****) respectively, all in the mid-late Miocene. The *bancana* group had its crown node dated at 8.83 mya (95% HPD: 7.06-10.68 mya).

The DEC model proved to be the most likely of the six ancestral area reconstruction models implemented in RASP (Supplementary material **Table S1**). The model clearly suggested Borneo as the most likely area of origin for the myrmecophytic clade (**Fig. 2**). Within the myrmecophytic clade, the crown nodes of all three myrmecophytic sections were also mapped with greatest probability to Borneo. Within section *Pachystemon*, only the *kingii* group seems to have exclusively evolved on Borneo. This island also came out as the most likely area of origin for the *hypoleuca* group but non-negligible probabilities were also assigned to wider distribution ranges of Malay Peninsula-Borneo (BE), Sumatra-Borneo (CE), and Sumatra-Malay Peninsula-Borneo (BCE). The *puncticulata* group seems to have a much broader region of origin with higher probabilities given to distribution ranges of BE and CE along with some support for BCE and Java-Borneo (DE). The MRCA of the *bancana* subgroup was also estimated to have had a wider distribution encompassing Borneo as well as Malay Peninsula (BE).

### 3.3. Ancestral state reconstruction

The most parsimonious reconstructions of the myrmecophytic state required six evolutionary changes for the Bayesian tree–one gain and five losses or two gains and four losses–and seven evolutionary changes for the RAxML tree–one gain and six losses or two gains and five losses (Supplementary material **Fig. S2**). The difference in the total number of evolutionary steps between the two trees can be directly attributed to the disparity in the resolution of the *bancana* group species (**Fig. 3**), notably of *M. rostrata* and *M. angulata*. Both cases, however, support one single instance of gain of myrmecophytism at the point of divergence of the myrmecophytic clade. This contradicts earlier hypotheses that assumed multiple independent acquisitions of domatia by the three myrmecophytic sections (Blattner *et al*., 2001; Davies *et al*., 2001). The single origin of myrmecophytism was apparently followed by multiple losses and possibly one secondary gain in *M. puncticulata*. Under the maximum likelihood method, the equal rates Mk1 model for ancestral state reconstruction (**Fig. 3**) could not be rejected with full confidence (p = 0.0546 for Bayesian tree, p = 0.07 for RAxML tree) when compared to the unequal rates AsymmMk model for both trees. Under the Mk1 model, the crown node of the myrmecophytic clade has a proportional likelihood of 0.85 to be a myrmecophyte in the Bayesian tree and 0.77 in the RAxML tree (**Fig. 3**). The reconstruction of the myrmecophytic state in the context of the *puncticulata* group is not clear under both parsimony and maximum likelihood methods, with its crown ancestor reconstructed as a myrmecophyte/non-myrmecophyte with equal (parsimony) or highly similar probabilities (maximum likelihood). The reconstruction suggested that the non-myrmecophytic state of *M. depressa*, *M. rostrata*, and *M. triloba* as most likely the result of a collective single loss for the Bayesian tree while for the ML tree, this was estimated as most likely to have involved two separate losses.

**Fig. 3.**
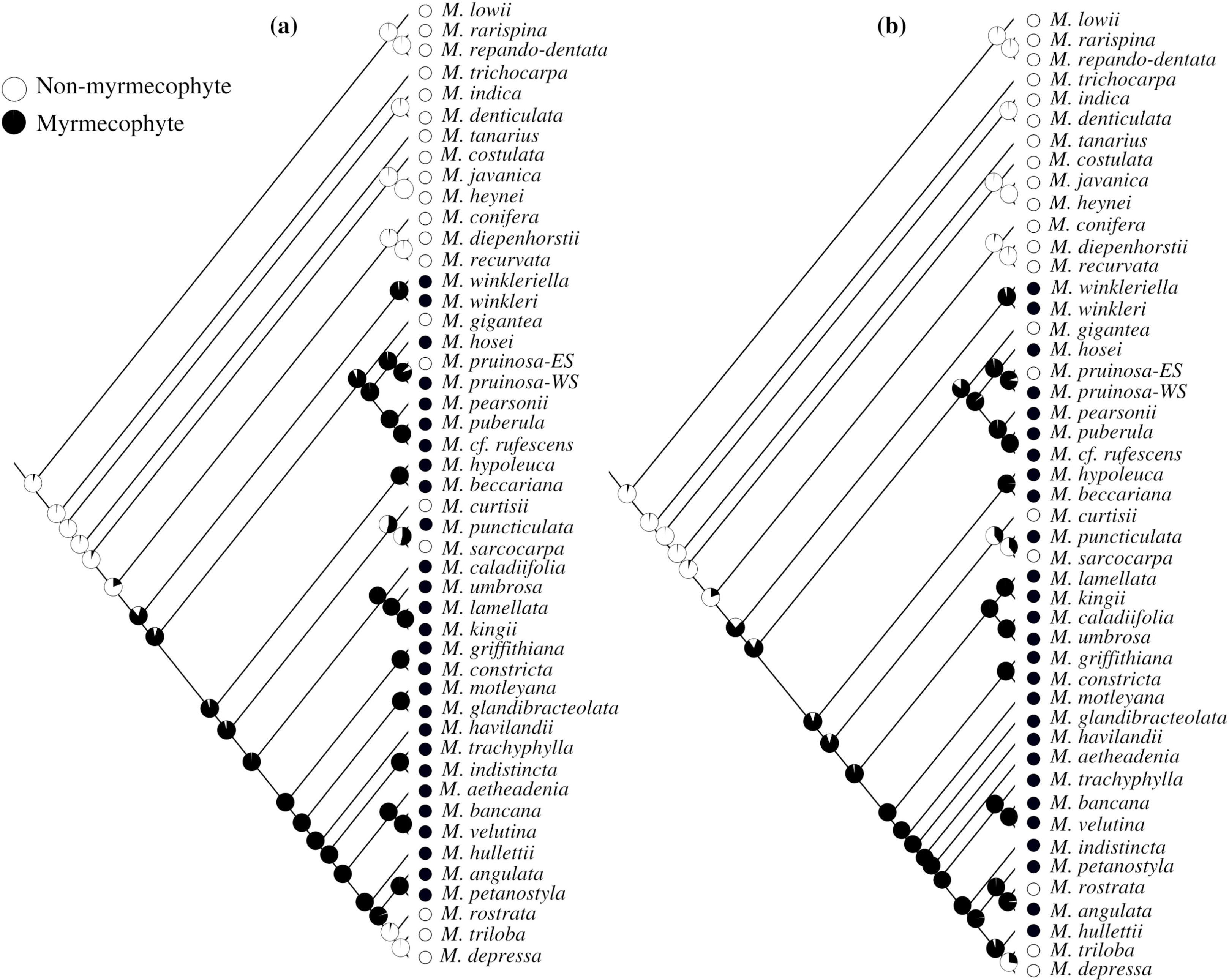
Ancestral state reconstruction of myrmecophytism (coded as a single trait) in *Macaranga* given by the Mk1 equal transition rates model implemented in the maximum likelihood framework on (a) BEAST2 maximum clade credibility Bayesian tree and (b) RAxML maximum likelihood tree. Pie charts at each node represent the probability of the ancestor being a myrmecophyte (black) or a non-myrmecophyte (white). Circles at the tree tips represent the present-day myrmecophytic state of the corresponding taxa.

### 3.4. Estimation of diversification rates

#### 3.4.1. Trait-dependent methods

For both the splitter and lumper treatments, BiSSE gave the strongest support to models that constrained the speciation and extinction rates to be equal for both states (A_0_=A_1_), indicating no differences in rates between myrmecophytes and non-myrmecophytes. The Bayesian posterior probability distributions of speciation rates showed an obvious overlap of their 95% credible intervals in both taxonomic treatments, again indicating no differences (**Fig 4a**).

**Fig. 4.**
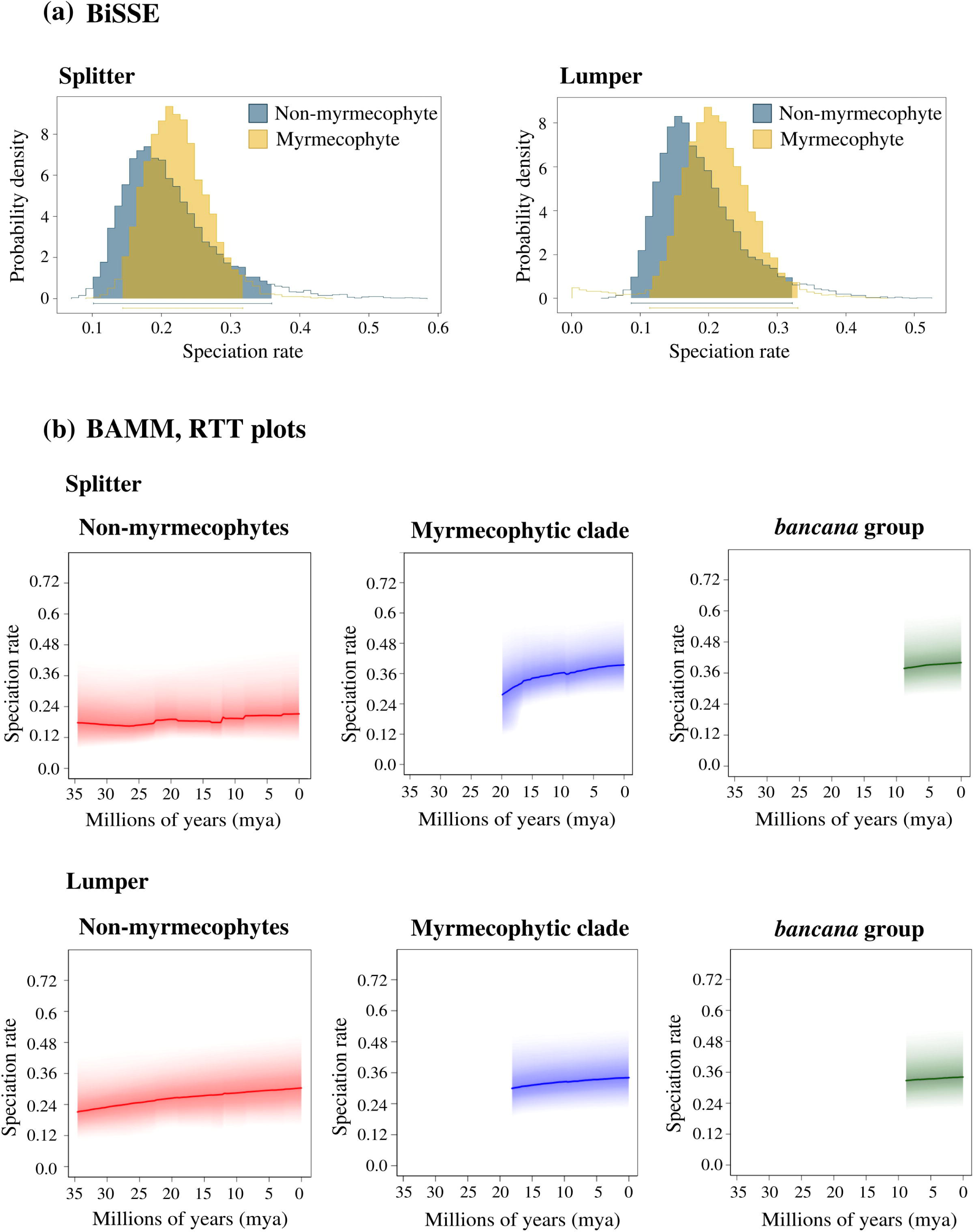
Diversification rate analyses estimates for myrmecophytic and non-myrmecophytic *Macaranga* (a) BiSSE estimates of posterior probability distributions of speciation rates for myrmecophytes and non-myrmecophytes under splitter (left) and lumper (right) treatments with prior values derived from the full BiSSE model estimation. Shaded areas and the horizontal bars below the distributions correspond to 95% credibility intervals. (b) BAMM diversification RTT plots for non-myrmecophytes (left), the myrmecophytic clade (centre) and the *bancana* group (right) for both splitter (top) and lumper (bottom) treatments. Colour shadings represent the 95% credible envelope on the distribution of rates at any point in time.

Under HiSSE, the results were less clear and also inconsistent between the two taxonomic treatments. In the splitter treatment, the full HiSSE model came out as the best model. Under this model, high speciation rate heterogeneity was detected but with myrmecophytes evolving much slower than the non-myrmecophytes (**Table 1**). The model also found one of the “hidden” states (A) diversifying faster when associated with non-myrmecophytes but not with myrmecophytes. In the lumper treatment under HiSSE, however, the null BiSSE model (A_0_=A_1_) was given the highest support, which suggests a lack of rate heterogeneity between the two states.

#### 3.4.2. Trait-independent method

Under the splitter treatment, BAMM detected a single rate shift with a posterior probability of 0.67 at ∼ 25 mya, well before the divergence of the myrmecophytic clade in *Macaranga* (Tabe **1b)**. It is interesting that this coincides temporally with the Asia-Australia collision which occurred between 25-23 mya. Through this tectonic event, the climate and vegetation of Southeast Asia underwent a drastic change resulting from the closing up of the Indonesian throughflow (de Bruyn *et al*., 2014). No rate heterogeneities, however, were detected under the lumper model as the zero rate shift model received the highest posterior support (**Table 1**). Clade-specific RTT plots for the splitter treatment showed slightly enhanced speciation rates within the myrmecophytic clade and within the *bancana* clade, although the difference does not appear significant (**Fig**. **4b**). Under the lumper treatment, the RTT plots showed a more stabilised speciation rate trend within the myrmecophytic and the *bancana* clade with no obvious differences in comparison to the non-myrmecophytic clade (**Fig**. **4b**). Overall, BAMM did not support a significant rate change associated with the divergence of the myrmecophytic clade ∼18 mya.

## 4. Discussion

### 4.1. Origins and evolutionary history of myrmecophytism in *Macaranga*: losses exceed gains

Our results indicate that myrmecophytism in *Macaranga* had an early-mid Miocene origin on the island of Borneo (**Fig**. **2**). Although the genus has a wide palaeotropic distribution spanning regions from West Africa to the islands of the South Pacific (Whitmore *et al*., 2008), the acquisition and retention of obligate ant-partners seems to be confined to a single Southeast Asian lineage in Sundaland. Many species across the entire distribution range in *Macaranga* participate in generalist ant-plant interactions and attract non-specific ant partners by providing EFNs and sometimes food bodies—traits that have been implicated as predispositions for the evolution of myrmecophytism (Whalen & Mackay, 1988; Fiala & Maschwitz, 1991, 1992b; Davies *et al*., 2001; Whitmore *et al*., 2008). In addition to these morphological and physiological predispositions, specific abiotic factors were probably necessary for such generalist interactions to transition into a more specialised mutualism. Aseasonality likely is one of the constraining factors for the successful acquisition and maintenance of myrmecophytism as it is a prerequisite for year-round food body production (Davies *et al*., 2001). In this context, Borneo’s long emergent history from at least the mid-Oligocene (Hall, 2012), persistence of everwet forest cover, and generally stable climate (Morley, 2012; de Bruyn *et al*., 2014) coupled with ant radiations in the tropical canopies in the Miocene (Brady *et al*., 2006; Moreau *et al*., 2006; Moreau & Bell, 2013; Chomicki & Renner, 2015) likely provided optimal conditions for myrmecophytism to evolve from generalist ancestors.

It has been previously hypothesised (Blattner *et al*., 2001; Bänfer *et al*., 2004, Weising *et al*., 2010) that sections *Pachystemon, Pruinosae,* and *Winklerianae* independently acquired the myrmecophytic trait, but our study suggests that this may not have been the case. Ancestral state reconstruction estimated a single instance of myrmecophytic origin for all extant myrmecophytic species (**Fig**. **3**). This disagreement could be owed to the lack of resolution of evolutionary relationships in previous investigations where monophyly of myrmecophytic *Macaranga* could not be unequivocally established. Besides this, the association of the three sections with distinct myrmecophytic traits, specifically, mode of formation of domatia and location of food body production (Supplementary material **Table S3**; Fiala & Maschwitz, 1992a, 1992b; Davies, 2001; Davies *et al*., 2001) has been put across as an argument in defence of independent acquisitions. However, cases of morphological distinctions can be observed within these sections as well (Supplementary material **Table S3**), and in the case of *M. pruinosa*, which is non-myrmecophytic on Borneo but myrmecophytic on Sumatra and Malay Peninsula (Davies, 2001), they occur within a single species. Cases of phenotypic plasticity in myrmecophytic traits, specifically domatia, have been reported in other genera such as *Tococa* where domatia production is influenced by water inundation of host plants (Izzo *et al*., 2018), and *Barteria* (Kokolo *et al*., 2020), which exhibits phenotypic plasticity in domatium size. Given these observations, it appears that myrmecophytic traits are generally plastic and the variation in traits among the *Macaranga* species could hence indicate divergent selection on plastic homologous traits by specific ecological factors and ant partners rather than independent acquisitions.

Four to six losses of domatia were inferred in *Macaranga* (**Fig**. **3**, Supplementary material **Fig**. **S2**), suggesting that losses have been more frequent than gains. Symbiotic mutualisms can break down and partners can revert to a free-living state if the cost-benefit ratio shifts so that costs outweigh benefits for one partner (Sachs & Simms, 2006; Chomicki & Renner, 2017). This is probably the case with *M. gigantea*, a pioneer species that can grow to massive heights (∼30 m) and known to grow exceptionally quickly under favourable soil conditions (Davies, 2001). It is likely that *M. gigantea* did not benefit much more from associations with obligate ant partners given that it can potentially outgrow its competitors and also employs abiotic chemical and physical defence mechanisms against herbivores (Nomura *et al*., 2000). Relying on an investment in these compensating features may have been economical for this species and the pay-off likely drove this branch to a loss of myrmecophytism. Historical biogeography estimated a wider distribution range that includes seasonal habitats such as Java (Heaney, 1991) for the crown ancestor of the *puncticulata* group (**Fig**. **2**). One could envisage a scenario where expansion of the distribution range to aseasonal habitats led to the loss of myrmecophytism in populations that left Sundaland and upon recolonizing retained the non-myrmecophytic trait. In this case, *M. puncticulata*, the only myrmecophyte in the *puncticulata* group, would represent a secondary gain. A similar argument has been proposed to explain the loss of myrmecophytism in the widely distributed *M. triloba* and Sundaland-restricted *M. depressa* by Davies *et al*. (2001). *M. rostrata*, a submontane species found at altitudes ranging from 800–2,300m (Davies, 2001) very likely represents a case of loss of myrmecophytism due to partner paucity at higher altitudes, a trend shown in other myrmecophytes as well owing to a decrease in ant species’ richness in higher elevations (Janzen, 1973; Bentley, 1977; Koptur, 1985; Longino *et al*., 2014; Gillette *et al*., 2015; Chomicki & Renner, 2017; Plowman *et al*., 2017).

### 4.2. Did myrmecophytism enhance diversification rates in *Macaranga*?

Various approaches such as HiSSE, BiSSE, and BAMM were employed to test if myrmecophytism enhanced diversification rates in *Macaranga*. We acknowledge that the number of tips in the tree was certainly a limitation in the implementation of these methods on our dataset (Davis *et al*., 2013; Kodandaramaiah & Murali, 2018). We nonetheless employed them to evidence any obvious asymmetry in diversification rates between myrmecophytes and non-myrmecophytes. The only significant rate heterogeneity was detected with HiSSE (splitter) which pointed to a decrease in diversification rates in the myrmecophytic clade (**Table 1a**, Supplementary material **Table S2**). The only indication of increased diversification rates in the myrmecophytic clade was detected with BAMM (splitter; **Fig**. **4b**), however, this did not appear significantly different from the non-myrmecophytic clade. The single rate shift supported in this case was estimated to have occurred much before the divergence of the myrmecophytic clade (**Table 1b**). Overall, it seems reasonable to state that there isn’t enough evidence from our data to suggest a considerable increase in diversification rates associated with domatia evolution (**Fig**. **4**, **Table 1**, Supplementary material **Table S2**).

Of the ∼160 origins of myrmecophytism in vascular plants known so far, most are associated with species-poor clades, although notable exceptions like Hydnophytinae and *Neonauclea* do exist (Chomicki & Renner, 2015). The myrmecophytic clade in *Macaranga* is also not species-poor with 29 odd species–including taxa that are genetically evidenced as morphotypes–but it is not possible to conclude with certainty that this relatively high species number is a direct consequence of the acquisition of myrmecophytism. Sections *Winklerianae*, *Pruinosae*, and the *bancana* group all have their crown nodes dated and more or less aligned at ∼ 9 mya in the late Miocene (9.64 mya (95% HPD: 4.83-14.43 mya); 8.95 mya (95% HPD: 8.03-9.49 mya); 8.83 mya (95% HPD: 7.06-10.68 mya) respectively, **Fig**. **2**). Given that their geographic origins all lie in Borneo (**Fig**. **2**), their divergence may be linked to historical geological events on the island rather than their mutualistic associations with ants. Volcanic activity and the rapid uplift of Mt. Kinabalu are reported to have occurred in northern Borneo in the late Miocene which coincides with these estimates (Hall, 2013; de Bruyn *et al*., 2014; O’Connell *et al*., 2018). This may have triggered diversification of lineages in *Macaranga* through niche partitioning, habitat fragmentation, and the opening up of novel habitats available for colonisation by ancestral species. Variation in geographical distributions and ecological requirements such as habitat, altitude, and soil preference is observed among species across the three clades (Supplementary material **Table S3**). In addition, the prevalence of endemic submontane species in northern Borneo, such as *M. puberula*, *M. petanostyla*, and *M. rostrata* (Supplementary material **Table S3**), corroborates this hypothesis.

### 4.3. Evolutionary patterns linked to degrees of specialisation in myrmecophytic *Macaranga*

Obligate ant-plant mutualism may in general hinder the development of high partner specificity owing to intense competition for hosts (Fiala & Maschwitz, 1990; Yu & Davidson, 1997; Feldhaar *et al*., 2000) and uncoupled reproduction and dispersal of plant and ant partners (Quek *et al*., 2004). Nevertheless, in myrmecophytic *Macaranga,* ecological observations (**Fig**. **1**, Supplementary material **Table S3**; Fiala *et al*., 1999) reveal varying degrees of ant-partner specificity, suggesting the prevalence of different host plant strategies in terms of specialisation. A case of extreme specialisation is observed in the two species of section *Winklerianae*, both of which associate with just one specific ant-partner species, *Crematogaster* msp8 (**Fig**. **1**; Fiala *et al*., 1999). Myrmecophytes of Section *Pruinosae* are mostly specialised on two ant partner species: *Crematogaster borneensis* and *C. linsenmairi* (**Fig**. **1**; Fiala et al., 1999). Members of section *Pachystemon*, however, seem to be the least specialised, with each species on average associating with four ant partner species (**Fig 1**; Fiala *et al*., 1999; Feldhaar *et al*., 2016). Various factors seem to determine partner choice from both the ant and plant sides. For example, the presence of a wax coating on the stems of host plants acts as an ant partner filter as only some *Crematogaster* ants are capable of moving on waxy surfaces (wax-runners, Federle *et al*., 1997; Feldhaar *et al*., 2010). Some species which grow in heath or peat swamp forests—*M. puncticulata*, *M. caladiifolia,* and occasionally *M. pruinosa* host generalist ant species in their domatia (Fiala *et al*., 1999; Davies *et al*., 2001). Besides this, ant partner preference is shown to vary with geographic location for some broadly distributed hosts (Fiala *et al*., 1999). What triggered extreme specialisation in some cases, however, is not clear. It has been suggested that highly specialised and obligate mutualisms where partners need each other for survival may be at a higher risk of going extinct due to fewer escape routes to other means of survival (Toby Kiers *et al*., 2010; Chomicki *et al*., 2019), especially in cases where partners have to associate every generation via horizontal transmission (Chomicki *et al*., 2019) as in the case of myrmecophytic associations. Mutualists that are not very specialised in the context of partner specificity could hence be considered more resilient (Toby Kiers *et al*., 2010). In this context, it is interesting to note that the more specialised *Winklerianae* and *Pruinosae* are much more species-poor compared to the less specialised *Pachystemon*, which may reflect higher extinction risks in the former sections.

### 4.4. Conclusion

We may conclude that myrmecophytism represents a delicate mutualistic system involving horizontal transmission of partners who are not able to survive without each other (Fiala *et al*., 1999). This fragility makes obligate mutualistic systems such as the *Macaranga-Crematogaster* symbiosis vulnerable to various environmental and ecological fluctuations (Briand *et al*., 1982; Fiala *et al*., 1999; Sachs & Simms, 2006; Toby Kiers *et al*., 2010). However, the degree of specialisation may determine the outcome of such disruptions. Species that have not become overly committed to a specialised partner may be more adaptable and able to survive such disruptions either through partner switches or even reversions to a free-living state if survival is mediated by other compensating traits (Bronstein *et al*., 2004; Moraes & Vasconcelos, 2009; Toby Kiers *et al*., 2010) or if defensive mutualisms are no longer economical. Although the evolution of myrmecophytism plays a significant role in the survivability of host plants in competitive habitats (Heil & McKey, 2003), it appears to be a rather opportunistic key innovation that is labile as long as it is not too specialised. Long-term stability of several environmental and ecological factors may lead some branches toward over-specialisation, in which case myrmecophytism might even represent an evolutionary dead end.

## Acknowledgements

We gratefully acknowledge the German Research Foundation (DFG, grant Gu1840/1) for funding this work. We thank Professor Kurt Weising for his helpful comments on the manuscript. We thank Maggie Bersch and Irene Diebel for helping out with the DNA extractions. The plant material used in our study was sourced from multiple field collection trips made to Southeast Asia from 1998 to 2014. We are grateful to the following people who took part in identifying and collecting plant leaf material on these trips: Christina Baier, Heike Feldhaar, the late Brigitte Fiala, J Jamsari, the late Ulrich Maschwitz, Ute Moog, Swee-Peck Quek, Ferry Slik, Natascha Wagner, and Malte Zirpel. We also thank Nurainas from the Herbarium of Universitas Andalas in Padang, Sumatra, for providing additional sample material. DG acknowledges the following for issuing collection permits: The Economic Planning Unit (EPU), Kuala Lumpur for collection in Sabah, Borneo; EPU in Kota Kinabalu, Sabah, and the Danum Valley Management Committee (2004/2005).

## CRediT authorship contribution statement

**Nadi M Dixit**: Conceptualization, Formal analysis, Investigation, Methodology, Visualization, Writing – original draft, Writing – review & editing. **Daniela Guicking**: Conceptualization, Funding acquisition, Investigation, Project administration, Supervision, Visualization, writing – review & editing.

## Data accessibility

Raw GBS sequencing data used in this study are deposited in the Sequence Read Archive (SRA) repository under the BioProject accession code: PRJNA1015024. The data will be available with the following link from 01.09.2024: https://www.ncbi.nlm.nih.gov/sra/PRJNA1015024

