## Supplementary material for "Myrmecophytism as a driver of macroevolutionary patterns: perspectives from the Southeast Asian *Macaranga* ant-plant symbiosis"

The following Supplementary Information is available for this article:

**Fig. S1** Maximum likelihood RAxML phylogenetic tree of 136 individuals representing 46 *Macaranga* species.

**Fig. S2** Most parsimonious ancestral state reconstructions of myrmecophytism in *Macaranga*.

**Fig. S3** STRUCTURE plot of 163 individuals representing 16 described species from the *bancana* subgroup of section *Pachystemon.*

**Table S1** Likelihood and AIC values for parametric biogeography models.

**Table S2** Parameter estimates for the best BiSSE and HiSSE models.

**Table S3** List of several ecological characters for all *Macaranga* species sampled in this study.

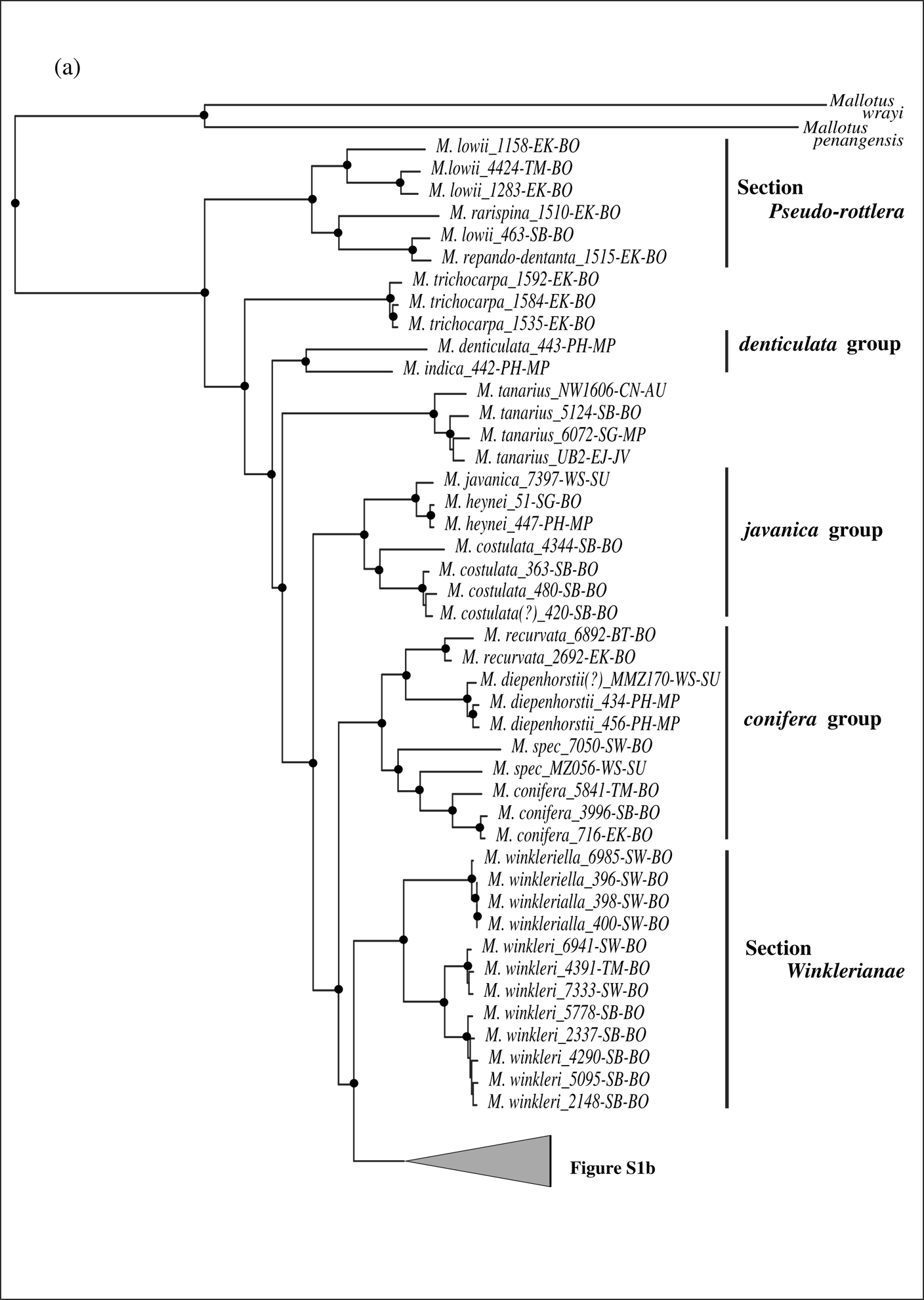

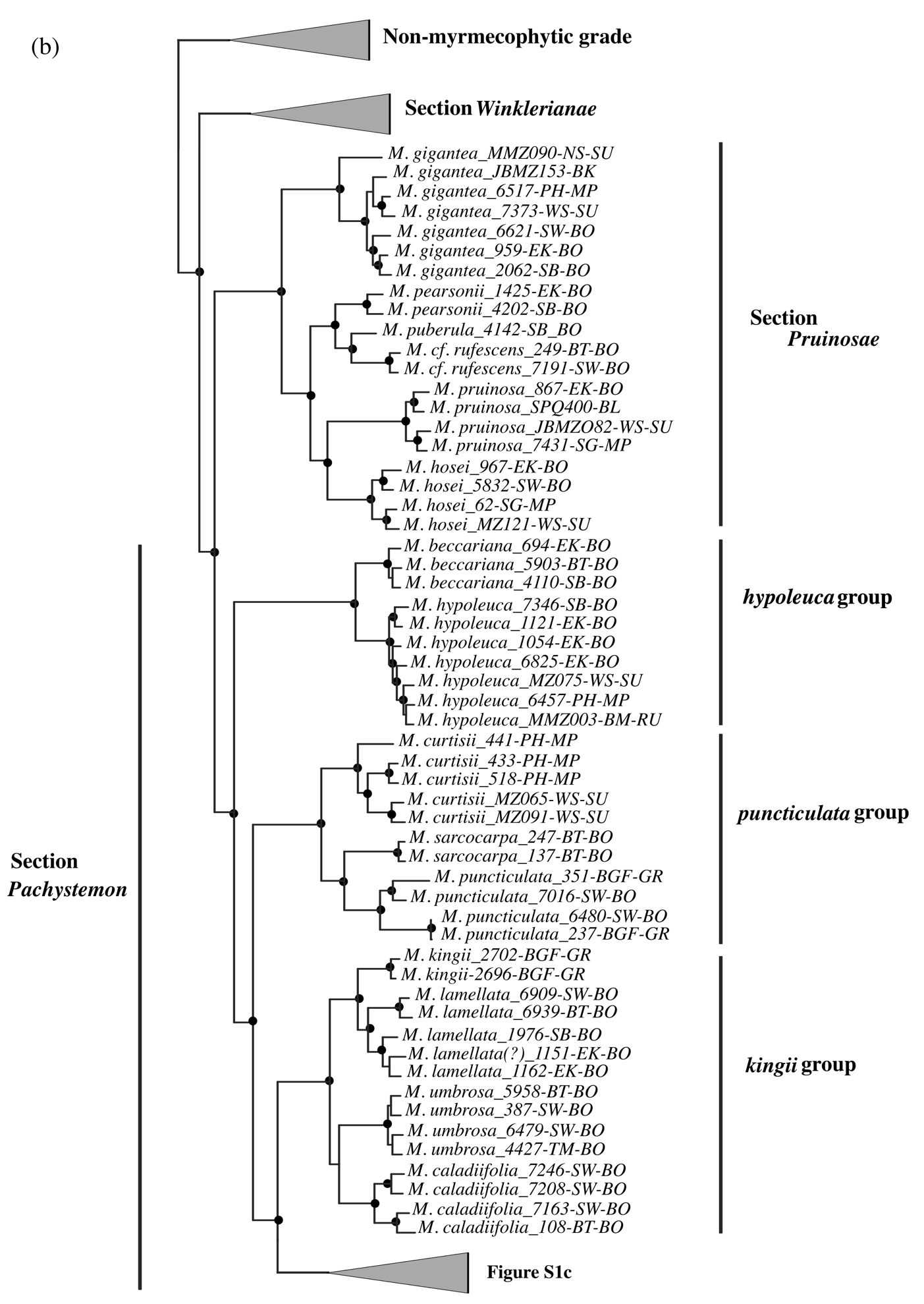

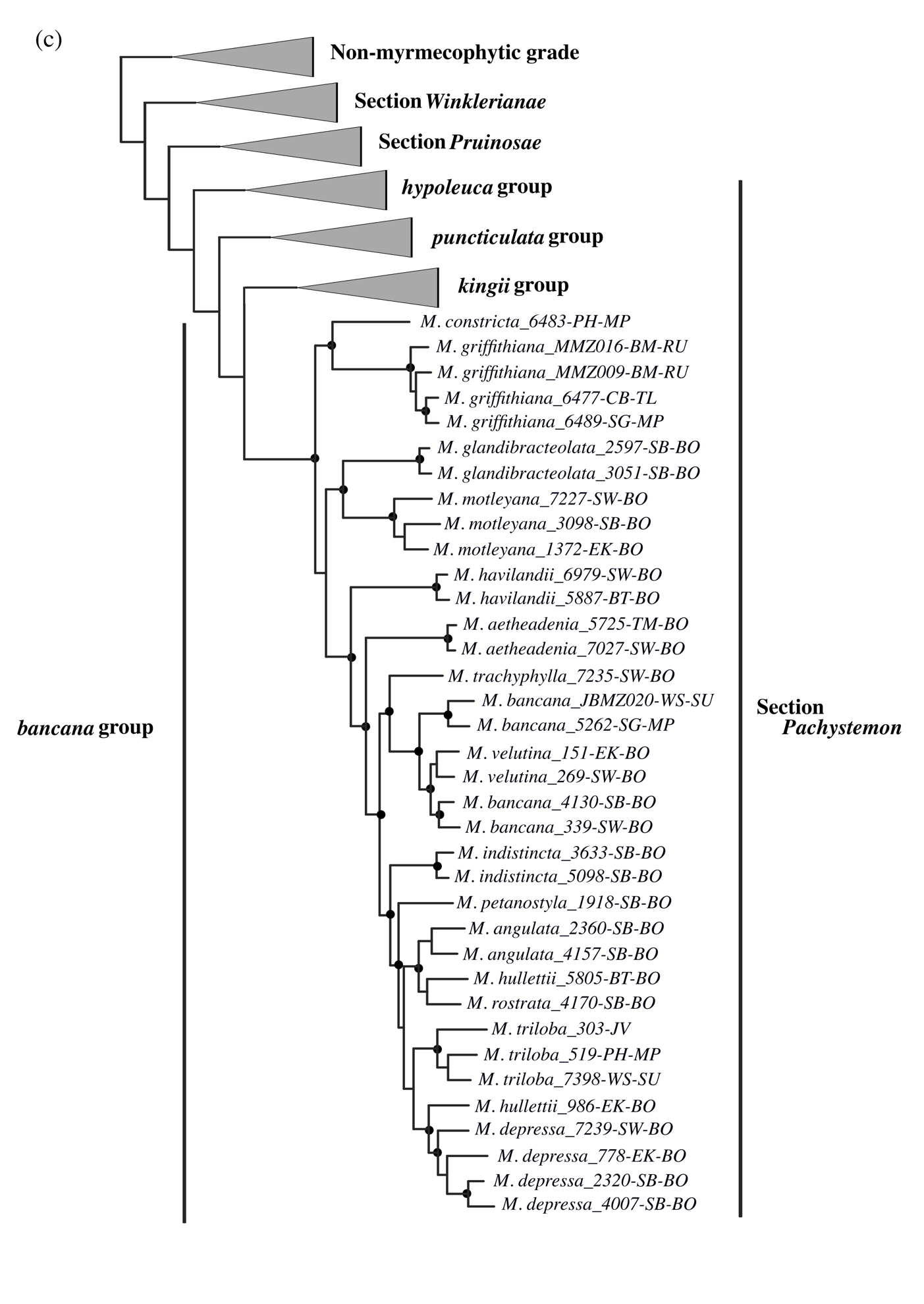

**Fig. S1** RAxML maximum likelihood tree estimation of 136 *Macaranga* individuals along with two *Mallotus* species functioning as outgroups. Sourcing of plant material, DNA extractions, GBS library preparations, and sequence alignment protocols to generate the sequence dataset used to calculate the tree followed Dixit *et al.* (2023), excepting two parameter settings in the ipyrad assembly pipeline: sequence clustering threshold parameter was set at 90% while the min_samples_locus parameter was set at 10% of the total number of samples - 14. The resulting sequence assembly had 1,279,038 sites for 138 individuals with 65.64% missing sites, representing a total of 10,071 loci. The SNP matrix had 146,508 sites with 58.90% missing sites. The tree calculation was performed using the ipa.raxml() tool available in the ipyrad analysis toolkit (Eaton and Overcast 2020). The tool automates the process of generating RAxML (Stamatakis 2014) command line strings and running them through Python code. For our data, the generalized time reversible (GTR) model was chosen as the substitution model along with the GAMMA model for rate heterogeneity (GTRGAMMA). A rapid bootstrap analysis with 100 replicates and the search for the best-scoring ML tree were simultaneously conducted in one single program run. The tree was rooted with the two *Mallotus* individuals: *Mallotus penangensis* and *M. wrayi*. Toytree (Eaton 2020) was then used to plot and visualise the best-scoring tree. Nodes that received bootstrap support > 90% are marked with black solid circles. Individuals are indicated in the following format: Species name_accession number-region of collection which are written as abbreviations: BGF, Botanical Garden Frankfurt; BK, Bangka; BL, Belitung; BM, Batam; BT, Belait; CB, Chanthaburi; CN, Cairns; EJ, East Java; EK, East Kalimantan; NS, North Sumatra; PH, Pahang; SB, Sabah; SG, Selangor; SW, Sarawak; TM, Temburong; WS, West Sumatra. Major regions of Australia, Borneo, Germany (only botanical garden collections), Java, Malay Peninsula, Sumatra and Thailand to which the aforementioned regions belong are abbreviated as AU, BO, GR, JV, MP, SU, and TL respectively. Individuals with a question mark (?) next to their species names or indicated as *M. spec* are those with uncertain species designations.

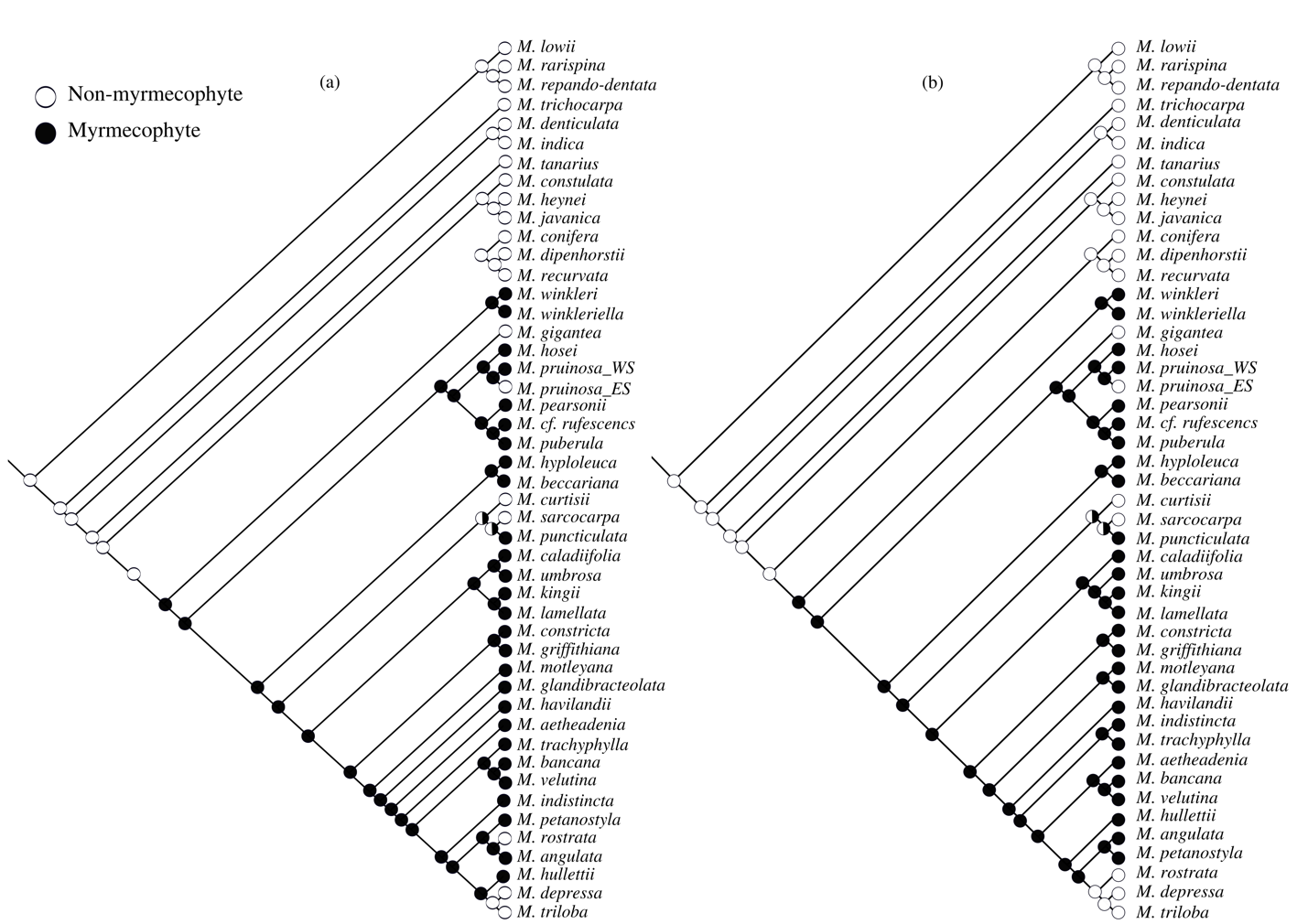

**Fig. S2** The most parsimonious ancestral state reconstructions of myrmecophytism in *Macaranga*, implemented on (a) RAxML maximum likelihood tree and (b) BEAST2 maximum clade credibility Bayesian tree with Mesquite (version 3.70; Maddison & Maddison, 2021). There were two most parsimonious reconstructions (MPRs) for each tree and the alternate parsimonious reconstructions are represented as equally likely states in the pie charts.

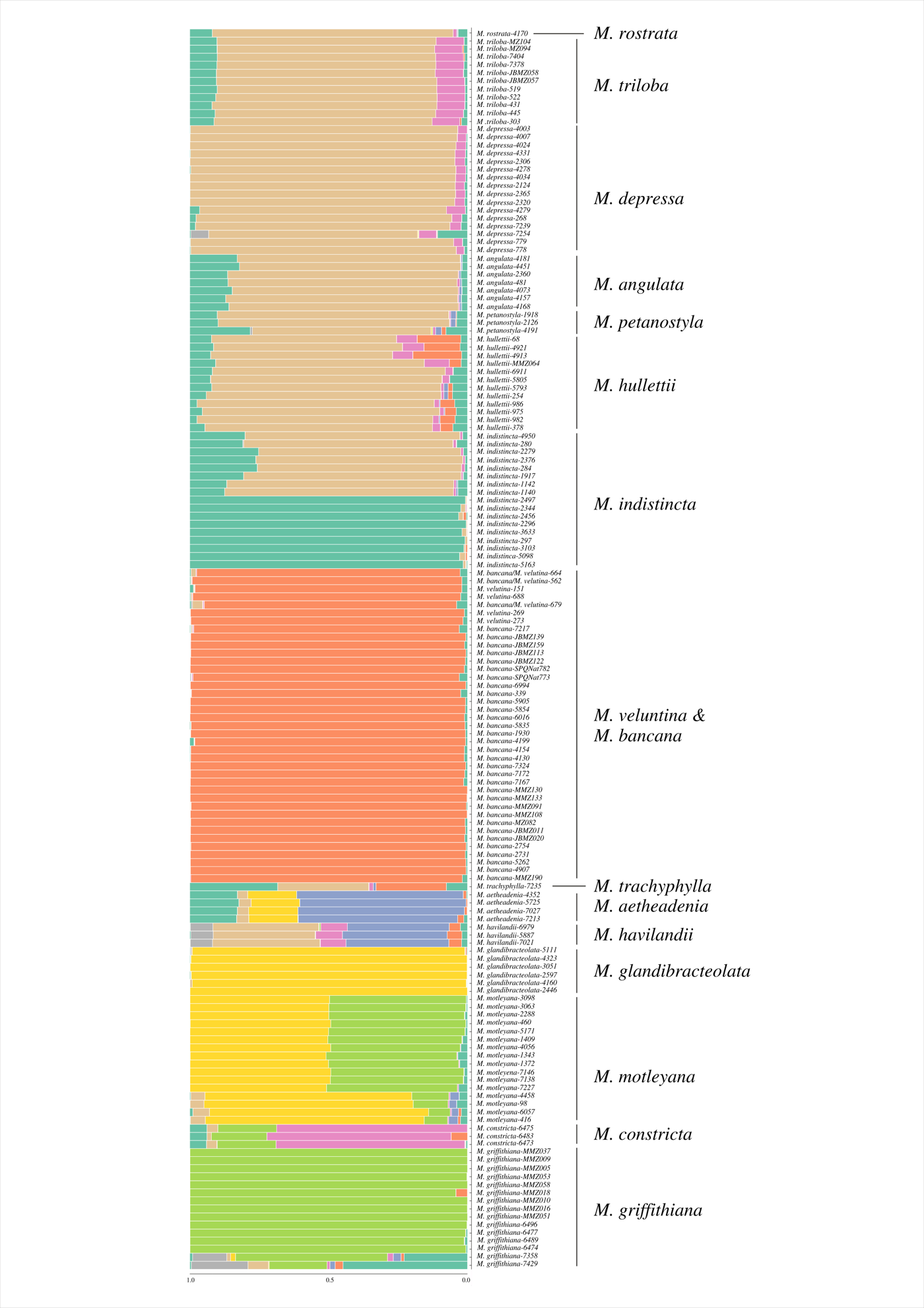

**Fig. S3** K= 9 STRUCTURE plot for 163 individuals representing 16 described species in the *bancana* group of section *Pachystemon*. The analysis was carried out with the STRUCTURE v.2.3.4 (Hubisz et al. 2009; Pritchard et al., 2010) tool available on the ipyrad analysis toolkit on GBS-derived SNP data. The admixture model was chosen as our ancestry model and the K parameter was allowed to vary from 3 to 20; each run was repeated 10 times. The rest of the parameters and priors were left at their default, recommended values (Pritchard et al., 2010). ΔK was calculated to identify the most probable number of clusters following the STRUCTURE run in accordance with Evanno et al. (2005). The calculation yielded greatest support to K = 9 genetic clusters.

**Table S1** Estimates of likelihood and AIC values for the six BioGeoBEARS ancestral area reconstruction implemented on RASP. The most supported model (*) is marked in bold.

| **Model** | **LnL** | **Number of parameters** | **d** | **e** | **j** | **AICc** | **AICc_wt** |
| --- | --- | --- | --- | --- | --- | --- | --- |
| **DEC *** | **-147.6** | **2** | **0.014** | **1.00E-12** | **0** | **299.5** | **0.66** |
| DEC + J | -147.6 | 3 | 0.014 | 1.00E-12 | 1.00E-05 | 301.8 | 0.21 |
| DIVALIKE | -149.5 | 2 | 0.016 | 1.00E-12 | 0 | 303.3 | 0.097 |
| DIVALIKE + J | -149.5 | 3 | 0.016 | 1.00E-12 | 1.00E-05 | 305.6 | 0.031 |
| BAYAREALIKE | -159.8 | 2 | 0.0092 | 0.037 | 0 | 323.9 | 3.30E-06 |
| BAYAREALIKE + J | -159.2 | 3 | 0.0092 | 0.03 | 0.0051 | 324.9 | 2.00E-06 |

**Table S2** Parameter estimates for the best BiSSE and HiSSE models under both lumper and splitter treatments. Under BiSSE, two states - 0 and 1- are specified to represent non-myrmecophytic and myrmecophytic states respectively. Under HiSSE, two hidden states - A and B - are specified in addition to ant-association states 0 and 1. Lambda, mu, and q represent speciation, extinction, and, transition rates respectively. q10 and q01 represent transition rates from a myrmecophytic state to a non-myrmecophytic state and vice-versa respectively. Additional transition rates (q0A1A, q0A0B, q1A0A, q1A1B, q0B0A, q0B1B, q1B1A, q1B0B) are specified under the full HiSSE model to indicate transition rates among the combined states 0A, 1A, 0B, and 1B. Transition rates for dual transitions involving both hidden and observed states (q0A1B, q1A0B, q0B1A, q1B0A) were dropped from all HiSSE models.

| Method | Best Model | state | lambda | mu | q10 | q01 | q0A1A | q0A0B | q1A0A | q1A1B | q0B0A | q0B1B | q1B1A | q1B0B |
| --- | --- | --- | --- | --- | --- | --- | --- | --- | --- | --- | --- | --- | --- | --- |
| BiSSE Splitter | lambda1 = lambda0, mu1 != mu0, q01 != q10 | 0 | 2.13E-01 | 1.02E-01 | 1.41E-01 | 3.81E-07 | NA | | | | | | | |
|  |  | 1 | 2.13E-01 | 5.87E-05 |  |  |  |  |  |  |  |  |  |  |
| BiSSE Lumper | lambda1 = lambda0, mu1 = mu0, q01 != q10 | 0 | 1.52E-01 | 3.71E-06 | 9.79E-02 | 6.11E-07 | NA | | | | | | | |
|  |  | 1 | 1.52E-01 | 3.71E-06 |  |  |  |  |  |  |  |  |  |  |
| HiSSE Splitter | Full HiSSE | 0A | 7.77E-01 | 5.15E-01 | NA | | 1.40E-01 | 2.06E-09 | 2.06E-09 | 4.91E-01 | 2.06E-09 | 2.06E-09 | 2.06E-09 | 4.92E-01 |
|  |  | 1A | 2.06E-09 | 5.15E-01 |  |  |  |  |  |  |  |  |  |  |
|  |  | 0B | 3.23E-01 | 5.15E-01 |  |  |  |  |  |  |  |  |  |  |
|  |  | 1B | 2.06E-09 | 5.15E-01 |  |  |  |  |  |  |  |  |  |  |
| HiSSE Lumper | Null BiSSE | 0 | 1.52E-01 | 2.06E-09 | 9.79E-02 | 2.06E-09 | NA | | | | | | | |
|  |  | 1 | 1.52E-01 | 2.06E-09 |  |  |  |  |  |  |  |  |  |  |

**Table S3**  A comprehensive list of several ecological characteristics of all *Macaranga* species sampled in the current study. Information on distribution, altitude, habitat and soil type was gathered from Davies (2001) and Whitmore *et al.* (2008); on ant partners from Fiala (1999) and Feldhaar *et al.* (2016); on domatia and its type from Fiala & Maschwitz (1992) and Davies *et al.* (2001); on food bodies from Davies (2001) and Davies *et al*. (2001); on wax cover from Davies (2001) and Federle *et al*. (1997); and on EFNs from Davies *et al.* (2001) and Fiala & Maschwitz (1991). (Table available as a separate spreadsheet).
